# Loss of Circadian Protection in Adults Exposed to Neonatal Hyperoxia

**DOI:** 10.1101/2020.07.10.197277

**Authors:** Yasmine Issah, Amruta Naik, Soon Y Tang, Kaitlyn Forrest, Thomas G Brooke, Nicholas Lahens, Katherine N. Theken, Amita Sehgal, George S. Worthen, Garret A. FitzGerald, Shaon Sengupta

## Abstract

Adverse early life exposures having a lasting negative impact on health. For examples, neonatal hyperoxia which is a risk factor for chronic lung disease of prematurity or bronchopulmonary dysplasia (BPD) confers susceptibility to respiratory infections like Influenza A (IAV) later in life. Given our previous findings that the circadian clock exerts a protective effect on injury from IAV, we asked if the long-term impact of neonatal hyperoxia includes disruption of circadian rhythms. We show here that neonatal hyperoxia abolishes the circadian clock mediated time of day protection from IAV, not through the regulation of viral burden, but through host tolerance pathways. We further discovered that that this dysregulation is mediated through the intrinsic clock in the lung, rather than through central or immune system clocks. Loss of circadian protein, *Bmal1*, in AT2 cells of the lung recapitulates the increased mortality, loss of temporal gating and other key features of hyperoxia-exposed animals. Taken together, our data suggest a novel role for the circadian clock in AT2 clock in mediating long-term effects of early life exposures to the lungs.

**Brief Summary:** Neonatal hyperoxia abrogates the circadian protection from Influenza infection in recovered adults.

## Introduction

Hyperoxia represents the single most important toxic exposure to premature neonatal lungs and is the key risk factor for chronic lung disease of prematurity or Bronchopulmonary Dysplasia (BPD)(1). Despite remarkable improvements in the survival of premature neonates, almost 40% of infants born at <29 weeks of gestation, (approximately 10,000 new cases in US each year) suffer from BPD(2, 3). In surviving adults, BPD is associated with an increased risk of respiratory infections, asthma(4, 5) and COPD(6–8). As an increasing number of prematurely born neonates survive into adulthood, these long-term morbidities have acquired greater public health importance(9). In a murine model, neonatal hyperoxia increased morbidity from Influenza A Virus (IAV)(10) in a dose dependent manner(11) through pathways involving both alveolar type 2 (AT2) epithelial cells and immune pathways(12).

We recently demonstrated that circadian rhythms offer a protection against IAV, wherein mortality is 3-fold lower if the animals are infected in the morning than in the evening(13). The mechanism involves both the pulmonary epithelium and the immune system. Circadian rhythms temporally segregate physiological process that allows organisms to anticipate its environment and adapt accordingly. The suprachiasmatic nucleus (SCN) is the master circadian pacemaker, however peripheral tissues, including the lung, also have cell autonomous clocks(14). Changes in oxygen tension are known to affect the clock(15, 16)as well as the interplay between the SCN and peripheral clocks(17). However, the effect of neonatal hyperoxia on circadian regulation of the recovered lung has not been investigated.

The early neonatal period represents a critical window for the development and consolidation of many important pathways, including circadian rhythms(18–20). Early life exposure to light(21, 22), inflammation(23) or alcohol has disruptive effects on circadian rhythms in adulthood(24). But, despite the high incidence of hyperoxia in premature neonates, how neonatal hyperoxia affects the development of circadian regulatory networks is not known. We hypothesized that early life hyperoxia disrupts the development of circadian rhythms and that such disruption undermines recovery from lung injury. Here we test this hypothesis using an IAV infection model of adult mice exposed to neonatal hyperoxia. We find that indeed exposure to neonatal hyperoxia abrogates the time of day protection from circadian regulation of IAV infection through primarily host tolerance rather than anti-viral effects. To identify the location of the relevant clock, we used tissue specific adult onset deletion of the core clock gene *Bmal1*. We report that disrupting *Bmal1* in AT2 cells of the lung faithfully recapitulates the phenotype of animals exposed to neonatal hyperoxia, suggesting that early life hyperoxia disrupts the circadian regulation of the pulmonary response to hyperoxia through the AT2 clock.

## Results

### Neonatal hyperoxia abrogates the circadian protection from influenza infection in recovered adults

We previously showed that mice infected at ZT11 (ZT11 refers to “Zeitgeber Time” 11 or dusk which marks the beginning of the active phase in mice since they are nocturnal) had three-fold higher mortality than mice infected at ZT23 (dawn or at the beginning of their rest phase)(13). Upon genetic disruption of the clock (*Bmal1^fl/fl^ERcre^+/-^*), in adult animals, the time of day difference was lost such that the mortality was high irrespective of the time of day of infection(15). Here we hypothesized that neonatal hyperoxia would disrupt circadian rhythms to result in a loss of temporal difference in outcomes in recovered adult mice infected with IAV at ZT11(dusk) or ZT23(dawn). Mice exposed to either hyperoxia or room air as neonates, were recovered to adulthood and infected with IAV (H1N1 mouse adapted strain PR8) at either ZT23 or ZT11 [Figure 1]. The room air controls maintained the time of day difference in mortality; the group infected at ZT11 had a mortality 2.6 times higher than the mice infected at ZT23 [Survival 72% in ZT23 vs 33% in ZT11, p <0.01]. By contrast, the mice that had been exposed to hyperoxia as neonates lost this time of day difference: both groups had mortality comparable to the ZT11 (in RA) group [Survival in hyperoxia groups: ZT23 at 37% and ZT11 at 25%; Figure 2 A and B]. This suggests that early life exposure to hyperoxia disrupts circadian regulation of the lung injury response to IAV infection.

**Figure 1.**
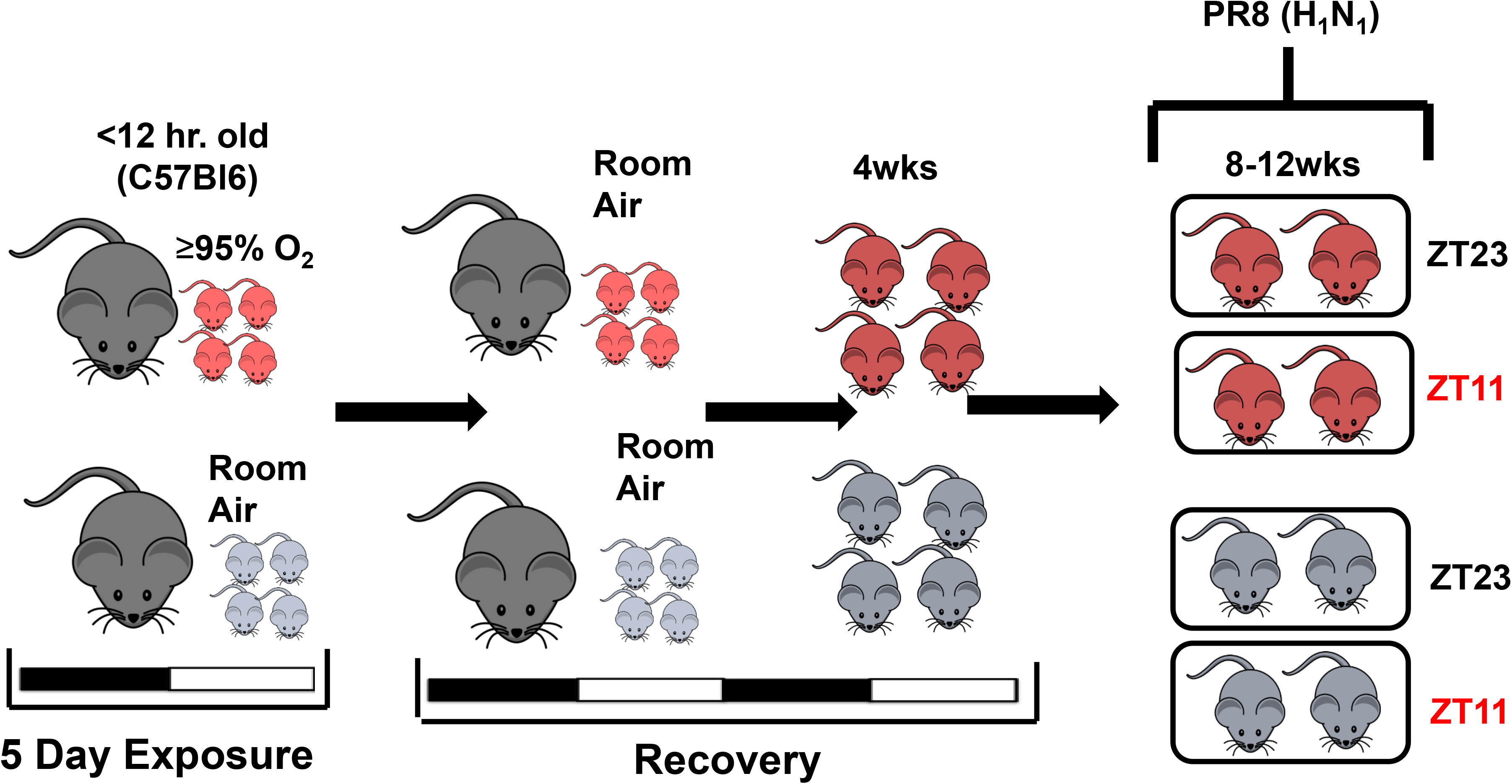

**Figure 2(A).**
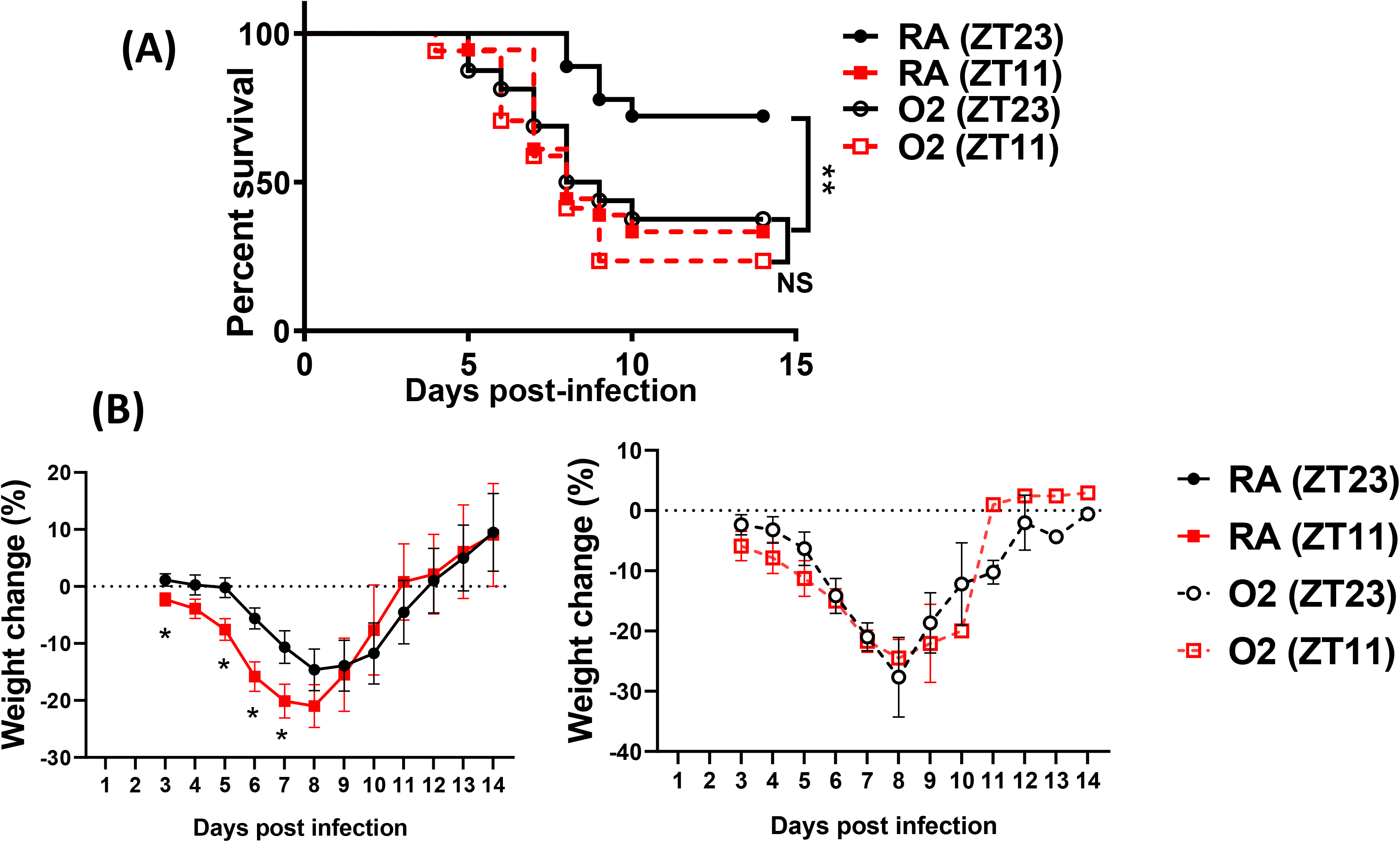

### Neonatal hyperoxia has only subtle effects on the central clock in adulthood

To determine the effect of neonatal hyperoxia on central circadian rhythms in adulthood, we evaluated the locomotor activity patterns of adult mice (8-12 weeks old of both genders) exposed to 5 days of neonatal hyperoxia (>= 95% O2 starting <12hrs of age to 5 days postnatal). A similar duration of hyperoxia is known to cause alveolar oversimplification, as seen in BPD(25). Immediately after exposure, the pups exposed to hyperoxia weigh less and appear sicker. However, these pups are indistinguishable from the room air controls in weight or overall activity by 8-12 weeks of age [Suppl. Figure 1]. To exclude differences in exercise tolerance, we measured locomotor activity using both infrared (IR) sensors and running wheels, and found that the two methods yielded similar results. Interestingly, while phase, period length, amplitude and total activity patterns were comparable between the two groups, the hyperoxia group displayed significantly more variation in their period lengths [Figure 3A].

**Figure 3(A).**
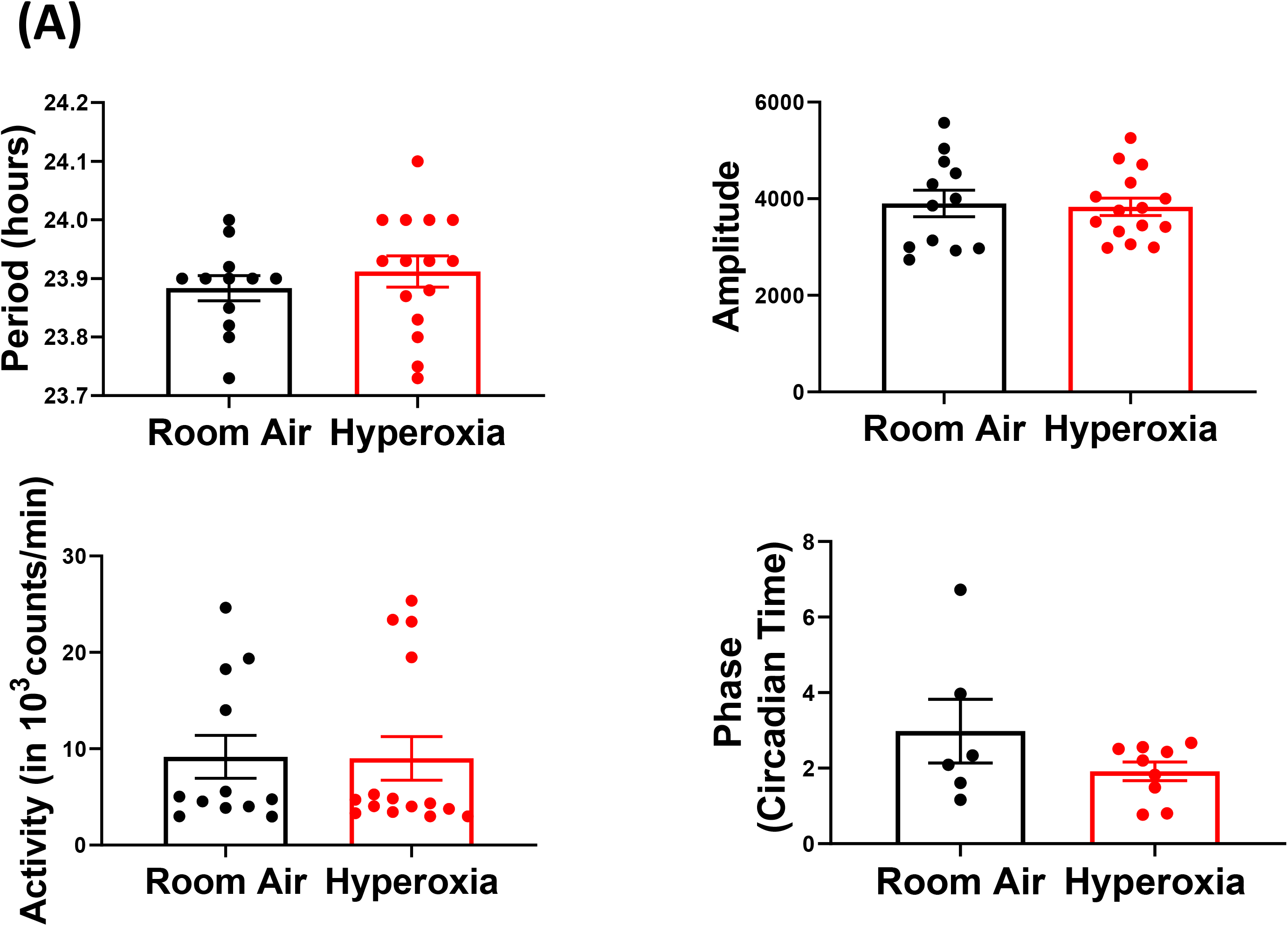

Interestingly, infection with IAV disrupts circadian locomotor rhythms under in a mouse model of COPD(26) raising the possibility that further stress may unmask a central phenotype. To address the hypothesis that neonatal hyperoxia caused instability of circadian regulation, we examined the ability of the animals to re-entrain to 12hr light-dark cycles after several weeks in constant darkness. Normoxia and hyperoxia groups did not entrain differently to this small change [Figure 3B and 3C]. Thus, considered together, these data are consistent with the hypothesis that effects of neonatal hyperoxia on IAV infection are not mediated via lasting changes in the central clock.

**Figure 3(B).**
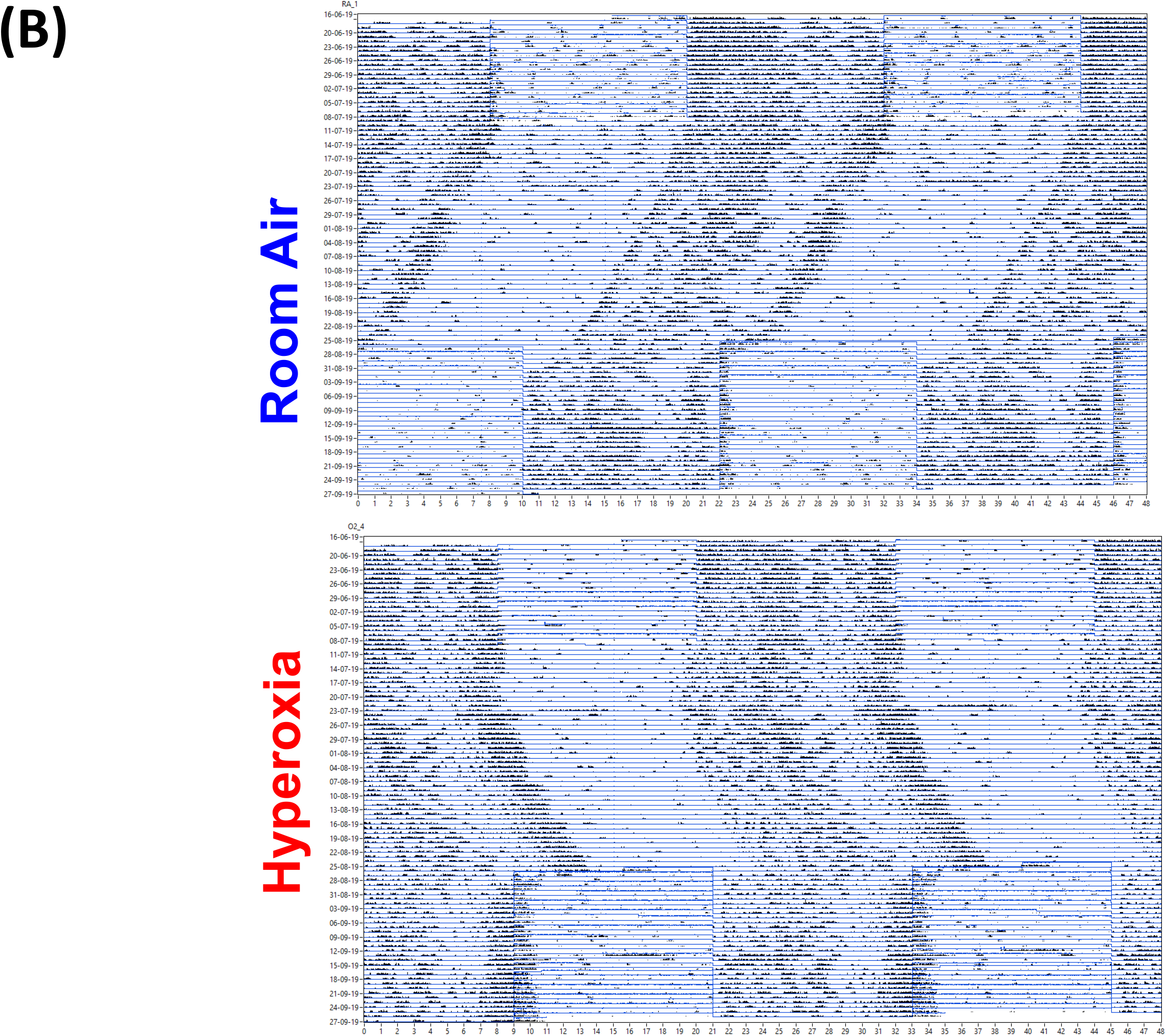

**Figure 3(C).**
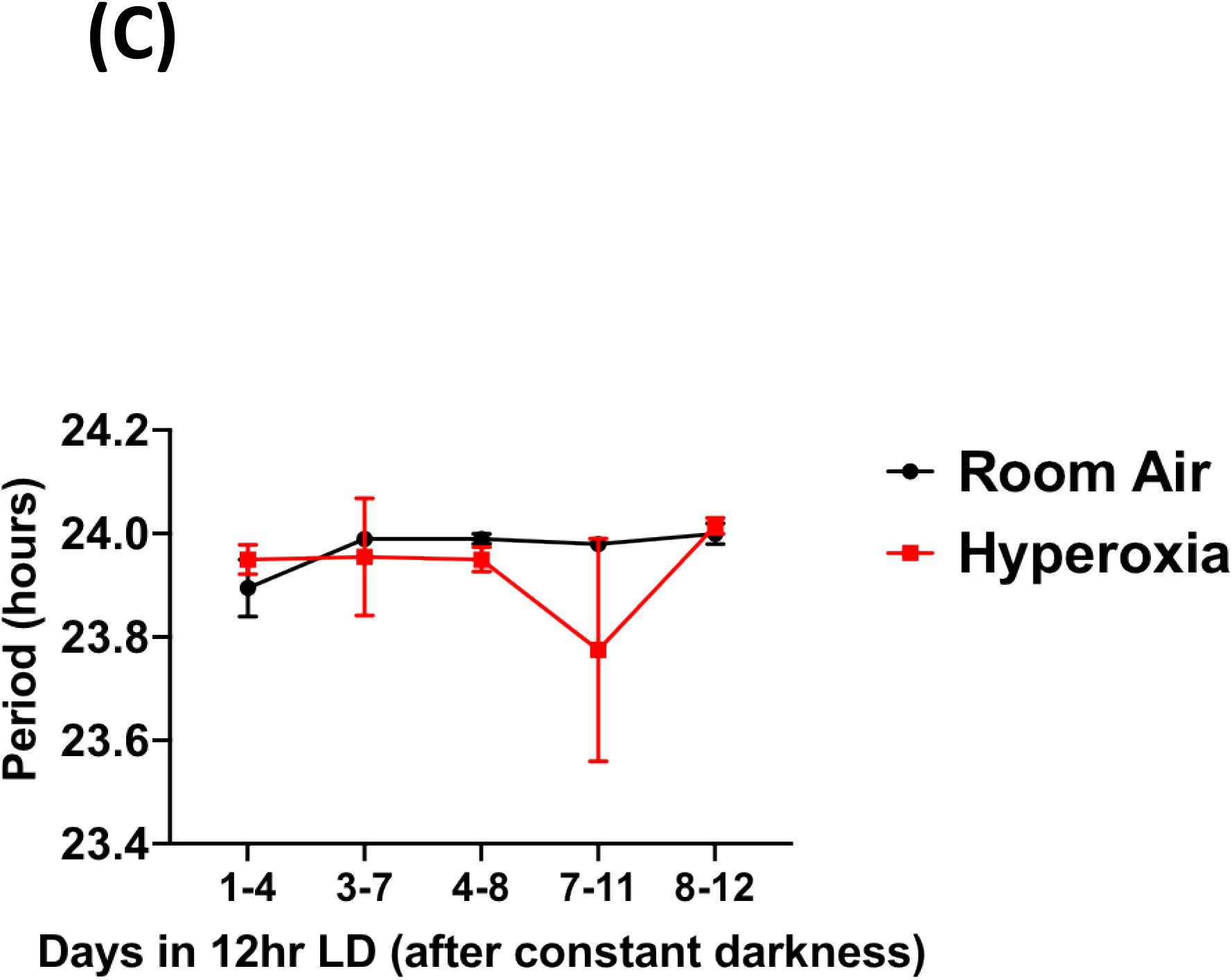

### Effect of neonatal hyperoxia on resident immune cells in the lung

We next addressed the possibility that loss of circadian gating of IAV responses in mice exposed to neonatal hyperoxia was secondary to circadian dysregulation of the immune response. We harvested lungs from adult animals exposed to either room air or hyperoxia as neonates and separated different populations of immune cells by flowcytometry at either the beginning of the rest phase (ZT23) or the beginning of the active phase (ZT11). While the total CD45^+^ population did not differ at these time points in the normoxia group, mice at ZT23 had higher numbers of CD45^+^ cells than at ZT11 in the hyperoxia group [Figure 4A]. The percentage of NK cells was significantly lower in the hyperoxia exposed mice at both time points. And the phase of the monocytes seems to reverse upon exposure to hyperoxia. Interestingly, in our previous work, these two populations, NK cells and inflammatory monocytes were different between the groups infected at ZT11 and ZT23(13). The fact that these two populations are perturbed in hyperoxia-treated animals implicates them in the circadian regulation of the IAV response.

**Figure 4(A).**
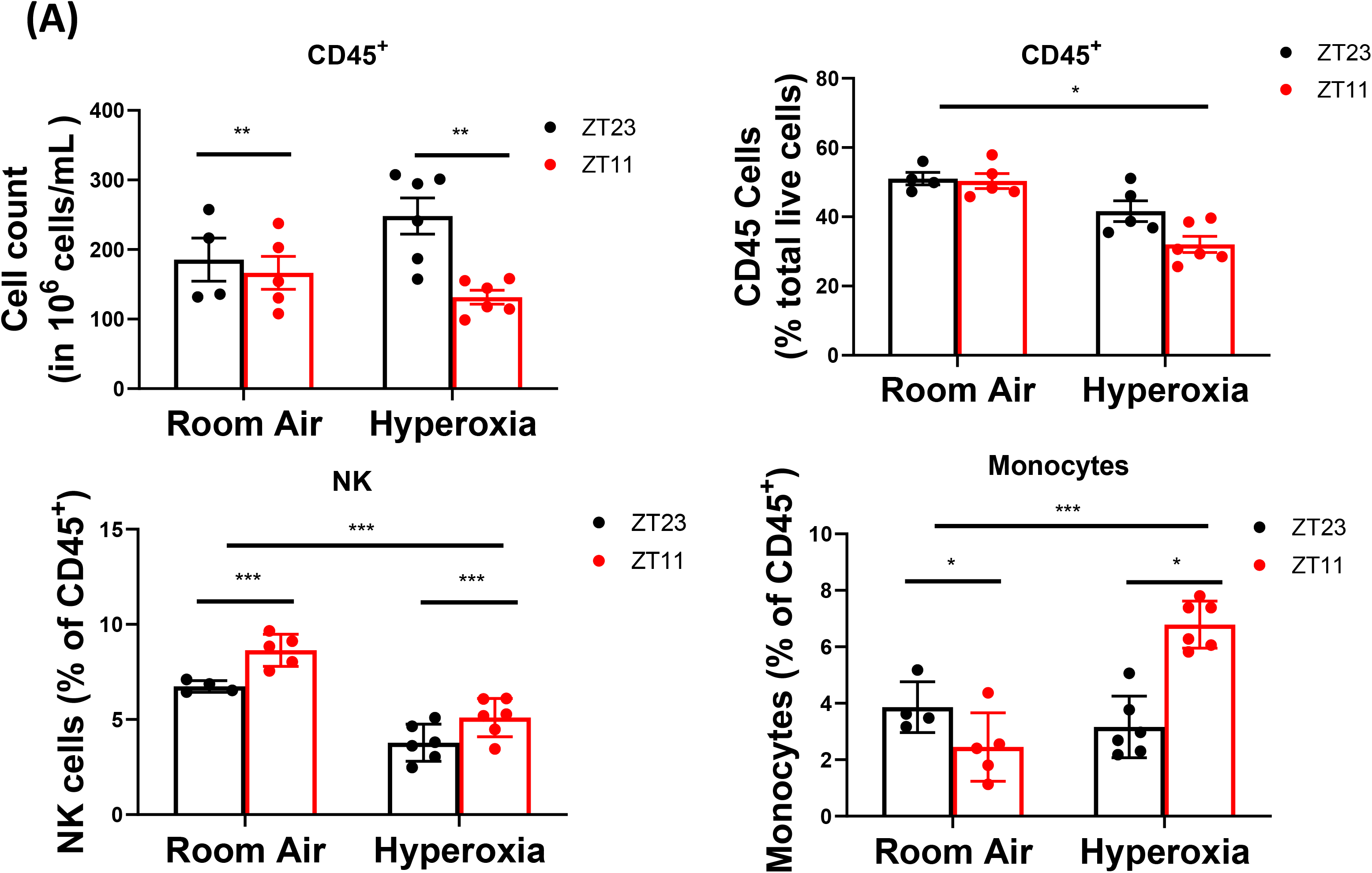

### Immune mechanisms underlying the circadian dysregulation of IAV induced lung injury in adults exposed to neonatal hyperoxia

We showed previously that circadian control of IAV induced lung injury and outcomes is not mediated by viral burden, but rather by the control of inflammation such that circadian mutants show exaggerated inflammation(13). Coupled with our findings above of impaired immune cell regulation in hyperoxia-treated animals, we hypothesized that the loss of time-of-day protection in these animals would be manifest as an exaggerated inflammatory response irrespective of the time of infection.

Consistent with our previous work, we found that the viral burden was comparable between the groups infected at ZT11 or ZT23; this held true in both neonatal hyperoxia and room air exposed groups [Figure 4(B]. However, on histological analyses, animals infected in the hyperoxia group scored worse for lung injury and a temporal effect was absent [Figure 4(C)]. To address the hypothesis that an exaggerated inflammatory response abrogated circadian variability in the innate immune response of animals exposed to neonatal hyperoxia, we performed bronchoalveolar lavage on days 1 and 5 following infection with IAV. Concordant with our previous work, among control animals exposed to room air as neonates, the room air group showed a higher total BAL count at ZT11 than at ZT23 by day 5p.i. [Figure 4(D)] among control animals exposed to room air as neonates. Contrary to our hypothesis, this was not influenced by neonatal exposure to hyperoxia, since even among the hyperoxia exposed pups, the total BAL count was higher in the ZT11 than in the ZT23 group. Thus, impaired circadian regulation of the response to IAV does not appear to arise from deficits in innate immune cells.

**Figure 4(B).**
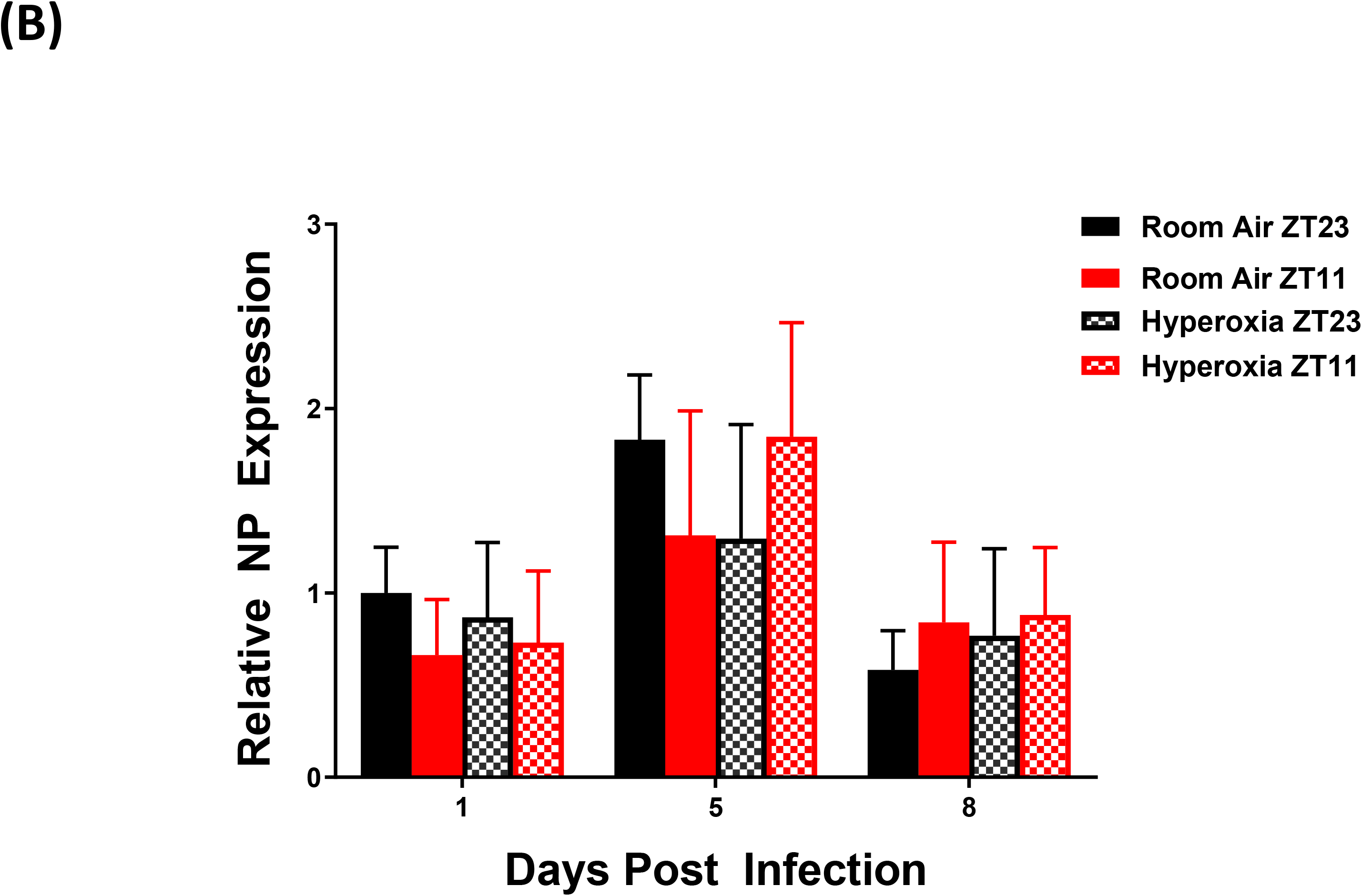

**Figure 4(C).**
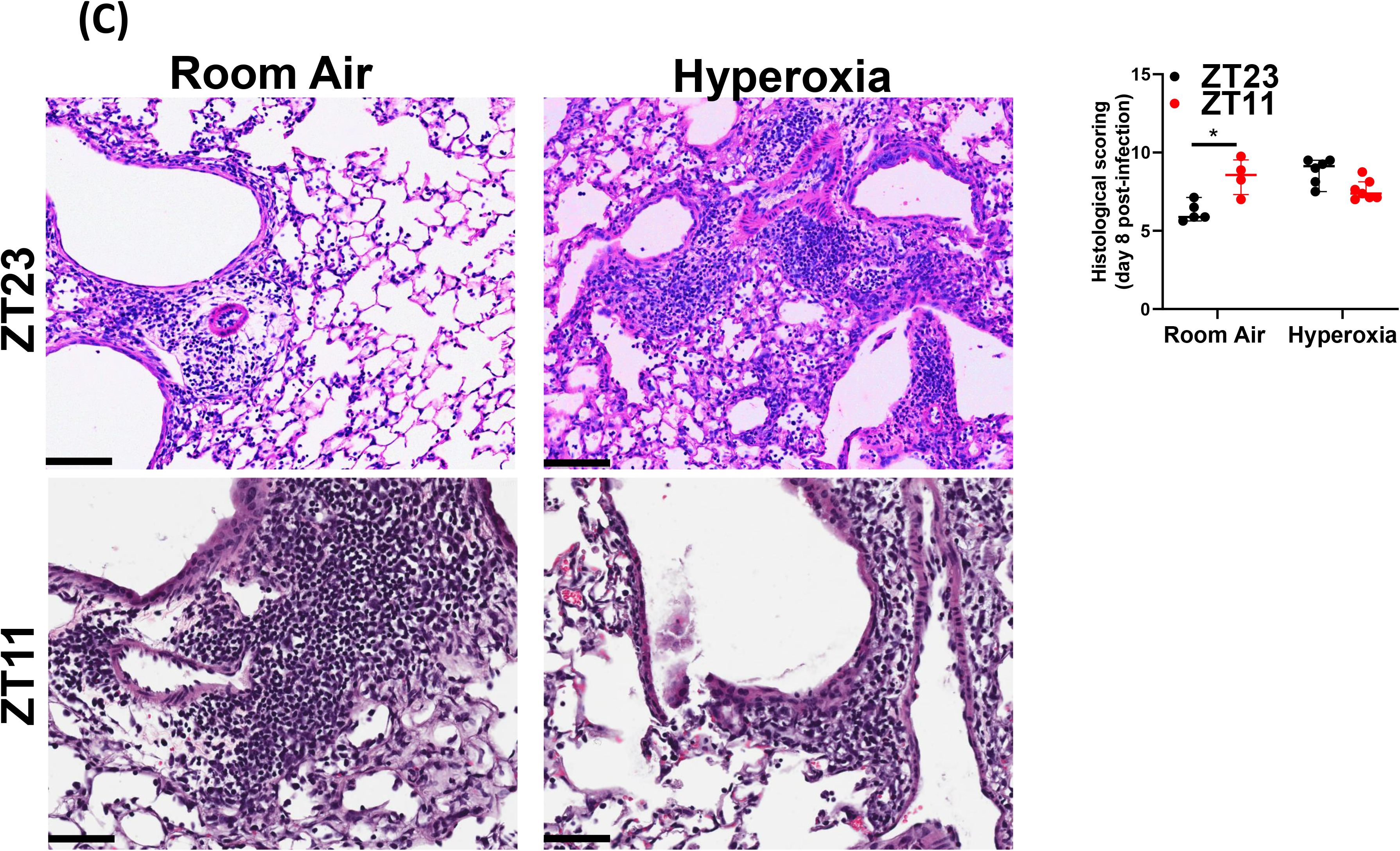

**Figure 4(D).**
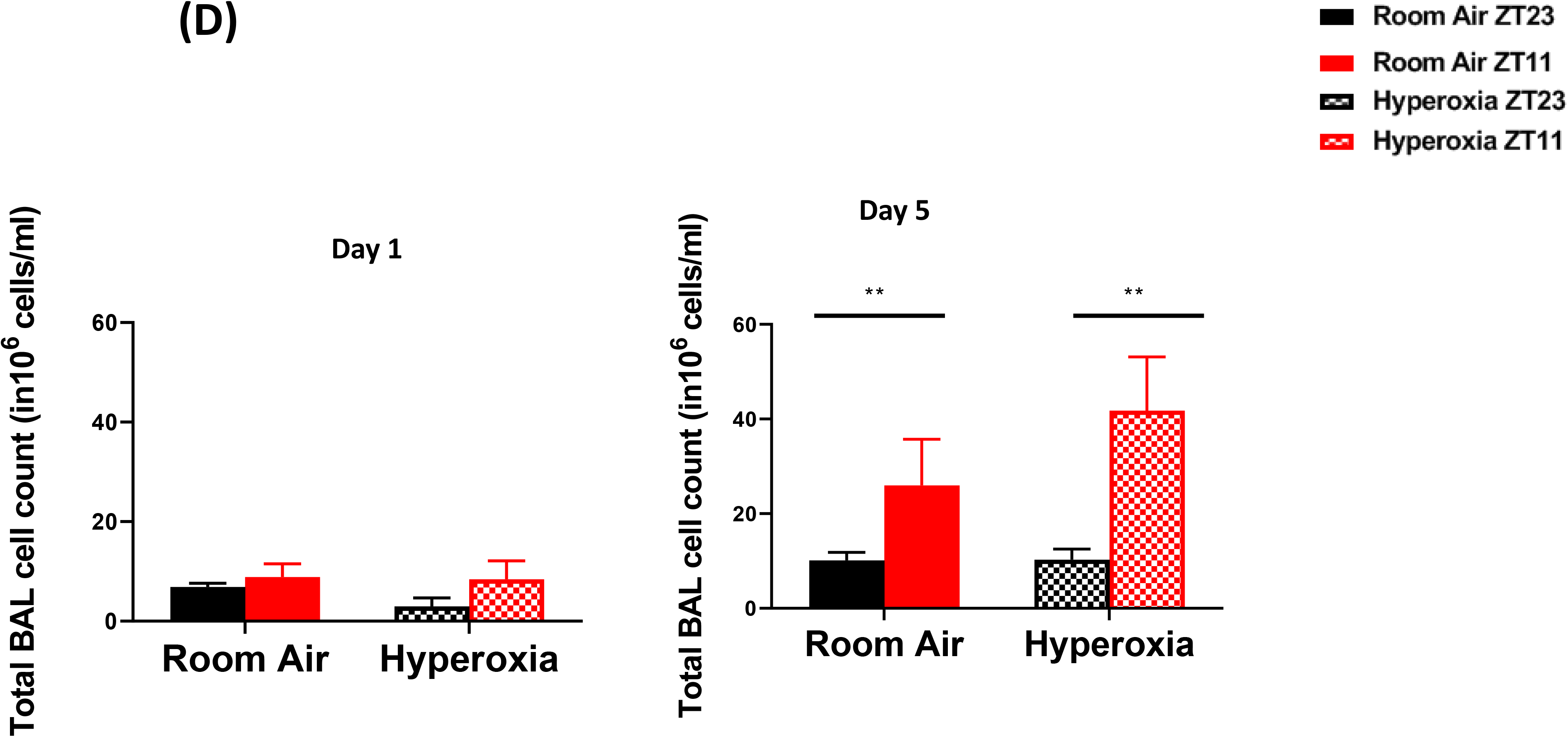

### Effect of neonatal hyperoxia on the pulmonary circadian system

We next investigated the effect of hyperoxia on the intrinsic clock in the lung by whole lung gene expression assays as well as with the use of a circadian reporter. While the phase of the oscillation of core clock genes *Bmal1, Per2 and nr1d2* was slightly altered by neonatal hyperoxia, the amplitude seemed intact. The amplitude of cycling of *Cry1* and Clock was slightly increased and the rhythm in *nr1d2* was apparently affected, but none of these changes were statistically significant [Figure 5A]. However, since bulk gene expression may conceal functionally relevant changes in cell specific circadian oscillation, we next used Per2luc mice, exposed them to hyperoxia as neonates and then recorded their bioluminescence *ex vivo* at 10-11 weeks of age. The hyperoxia exposed animals showed rhythmic expression of the Period2 gene with comparable periodicity and phase to the animals exposed to room air. However, the amplitude of these oscillations was significantly dampened in the hyperoxia group [Figure 5 B]. These data are consistent with the hypothesis that the perturbation induced by IAV infection unmasks a dysregulation of circadian function intrinsic to the lung in animals exposed to neonatal hyperoxia.

**Figure 5(A).**
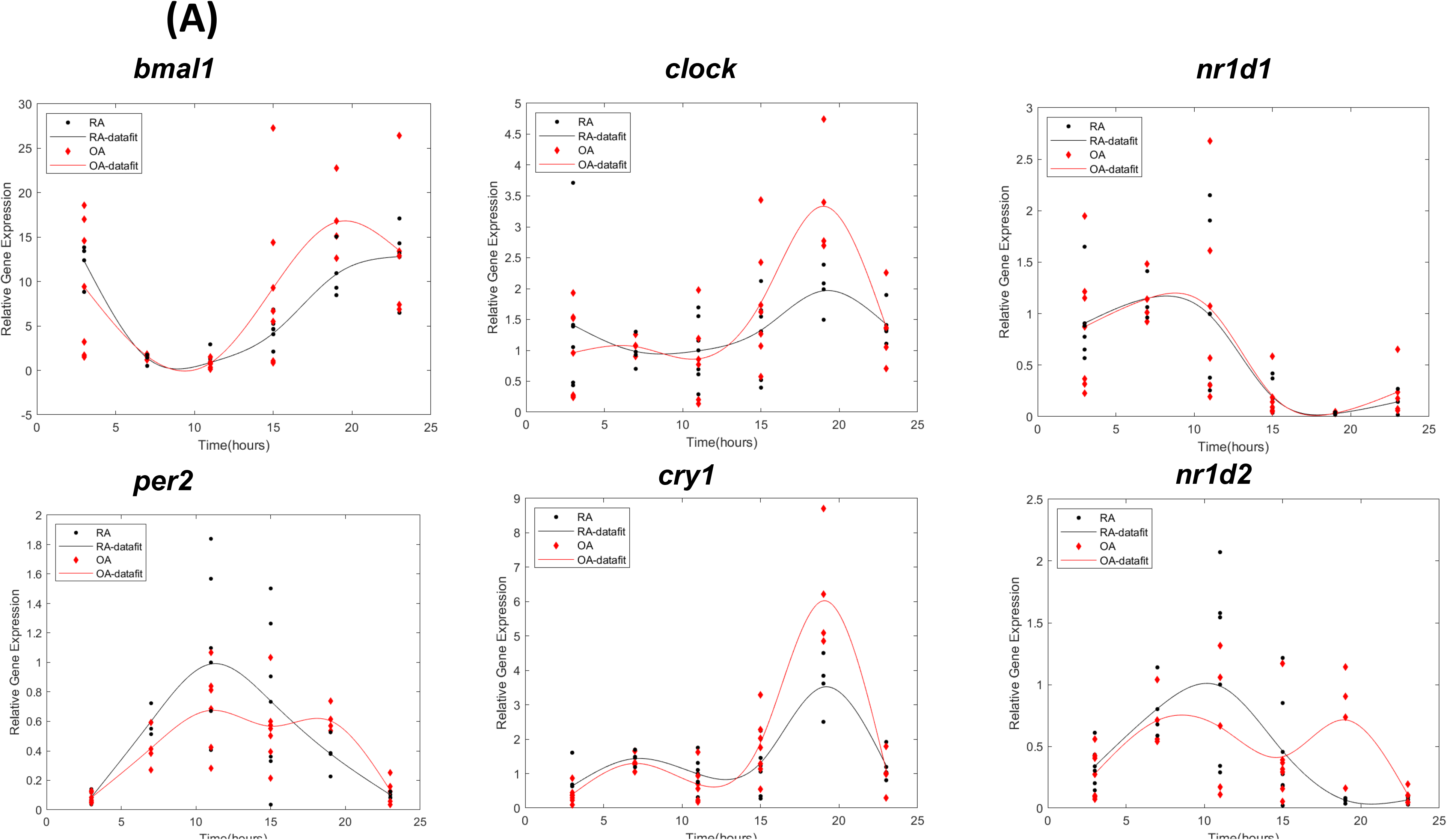

**Figure 5(B).**
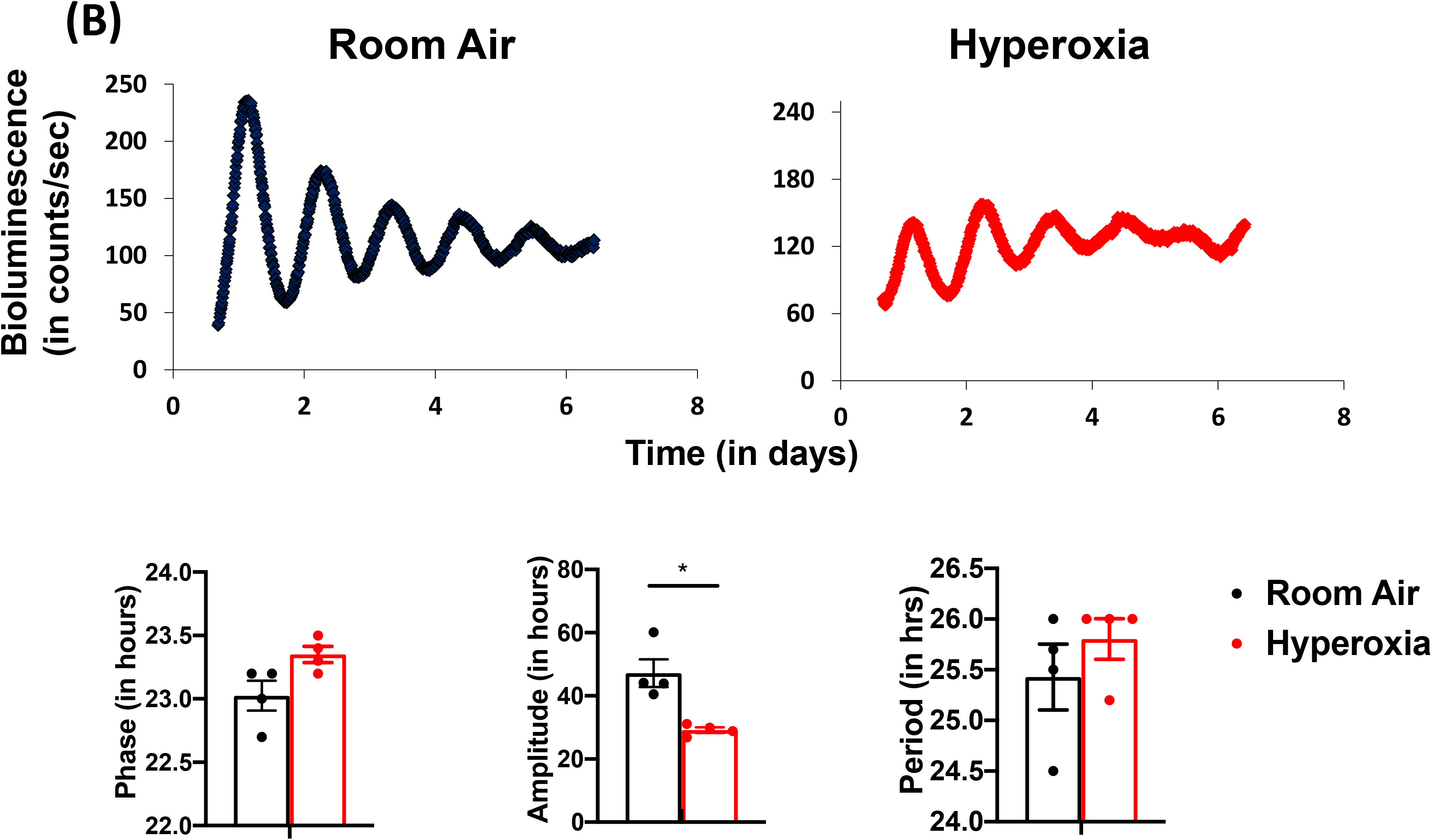

### Disrupting the circadian clock in AT2 cells in adulthood recapitulates the loss of time of day protection seen in adults exposed to neonatal hyperoxia

Neonatal hyperoxia disrupts the development of the respiratory epithelium, at least in part due to the depletion of AT2 cells after recovery(27). During hyperoxia, the AT2 cell number expands as a source of AT1 cells; upon return to room air for recovery, there is a progressive depletion of AT2 cells, with approximately 70% surviving at 8 weeks of age(27, 28). Thus, given the role of the type 2 airway epithelial cells in hyperoxic injury and repair, we asked clock disruption in AT2 cells would be sufficient to re-capitulate the effects of neonatal hyperoxia in terms of loss of circadian gating in response to IAV. An effect localized to AT2 cells would also be consistent with the relatively intact circadian gene expression in hyperoxia-treated mice, given that AT2 contribute only a small percentage of the cells in the total lung [Figure 5 A].

To test the relevance of the clock in AT2 cells, we generated mice in which the core clock gene, *Bmal1* was deleted in these cells [*Bmal1^fl/fl^Sfptc-Ert2cre^+^*]. While the cre-mice infected in the morning had better survival than cre-mice infected in the evening [Figure 6(A) 72% survival in CT23 group versus 20% survival is CT11 group; p<0.01 by Mental-Cox test], this time of day difference was lost in mice lacking *Bmal1* in their AT2 cells (25% in the CT23 group versus 38% in the CT11 group of *SfptcCre^+^Bmal1^fl/fl^;* CT refers to the time corresponding to ZT11 in constant darkness; In models of *Bmal1* deletion, we and others have used constant darkness conditions to avoid any differential effects of light on various peripheral clocks(13, 29, 30)). Post-natal deletion of *Bmal1* specifically in AT2 epithelial cells thus recapitulated the impact of neonatal hyperoxia on IAV induced mortality and morbidity.

**Figure 6.**
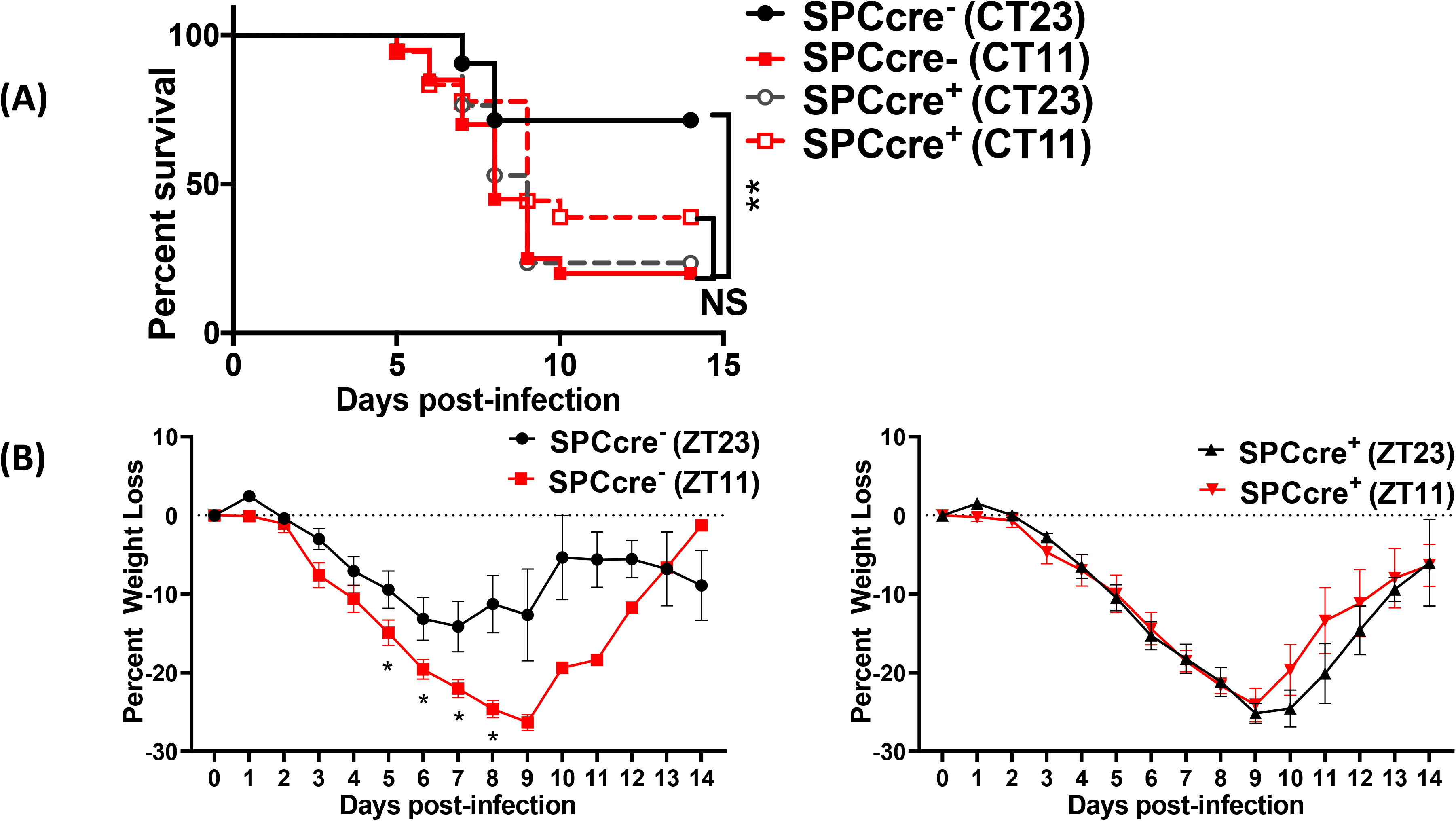

### *Bmal1* deletion in AT 2 cells reiterates the phenotype of circadian dysregulation seen in adult animals exposed to hyperoxia as neonates

As loss of circadian protection may be caused by different mechanisms, we sought to determine whether this loss in the AT2-Bma1^-/-^ animals was associated with similar underlying mechanisms as those one seen in hyperoxia exposed animals. Specifically, we determined whether the loss of circadian protection was mediated through anti-viral or host tolerance pathways.

We found that the viral loads at day 5 and day 8 were comparable across genotypes in animals infected at ZT11 or ZT23. This result is identical to that seen in hyperoxia treated animals exposed to IAV [Figure 6D]. We next compared the histology of the cre^+^ and the cre^−^ animals infected with IAV and found, again, that it was reminiscent of the hyperoxia treated animals. While the lung pathology was worse in cre-group infected at ZT11 than at ZT23, this time of day protection was lost in the cre^+^ animals, where the histology was worse at both ZT11 and ZT23. Further, we found that many of the gene expression signatures were comparable between the hyperoxia treated animals and those with Bmal1 deleted in AT2 cells. This included a reduction in *Il1b* and *il10* in both hyperoxia exposed animals as well as cre^+^ mice, while the pattern of change was comparable for *ccl2* [all on day 5 p.i. in Figure 6E]. Finally, even with the loss of *Bmal1* in AT2 cells (*Bmal1^fl/fl^Sfptc-Ert2cre^+^*), the BAL cell count varied by time of day [Figure 6F], being higher in ZT11 than at ZT23 in both cre+ and cre-animals on day 5 p.i.— again similar to the BAL count patterns seen in adults exposed to hyperoxia. Considered together, these results support the possibility that early life hyperoxia disrupts the circadian rhythm mediated protection in IAV induced lung injury through AT2-clock mediated effects.

**Figure 6(C).**
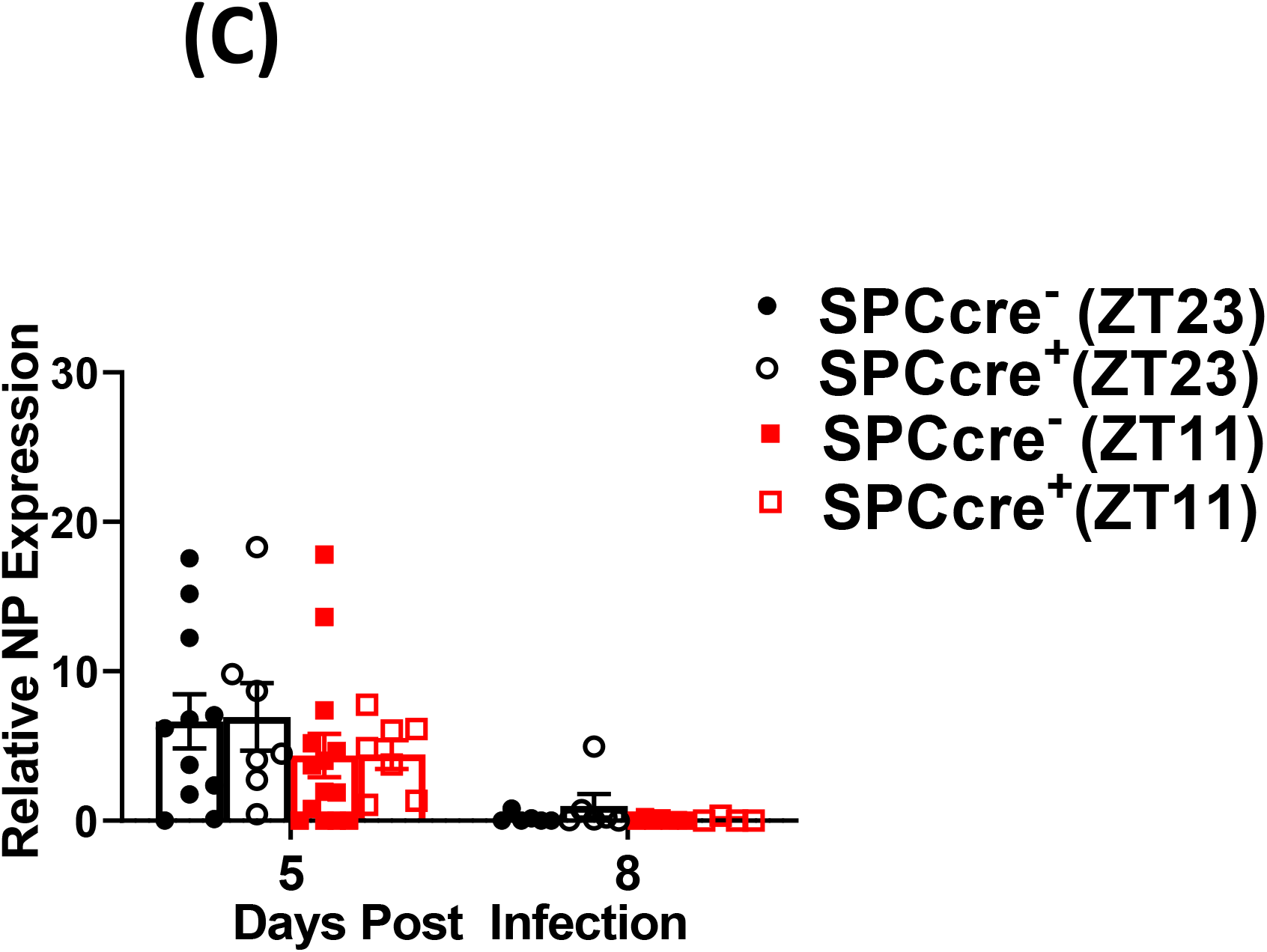

**Figure 6(D).**
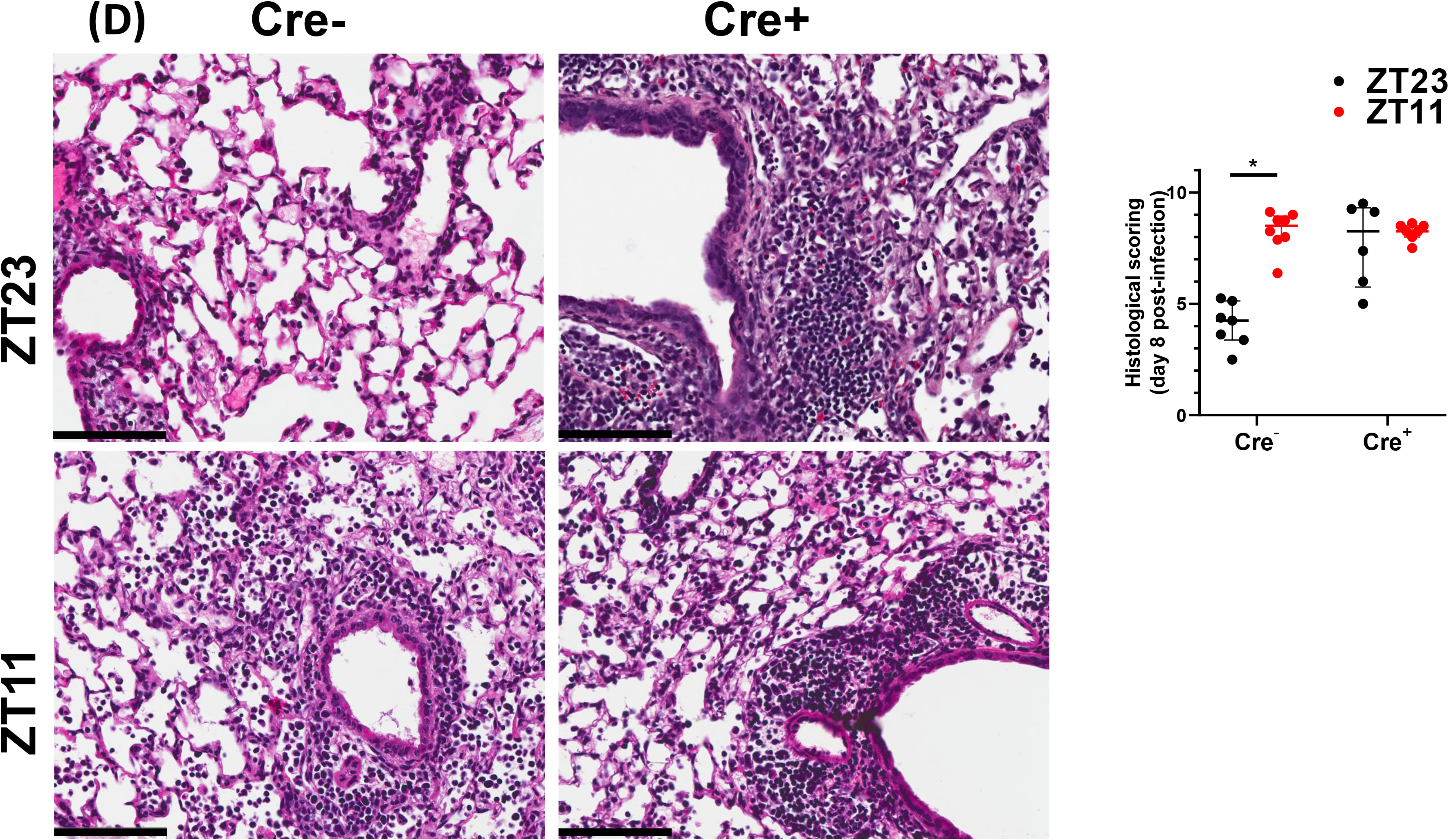

**Figure 6(E).**
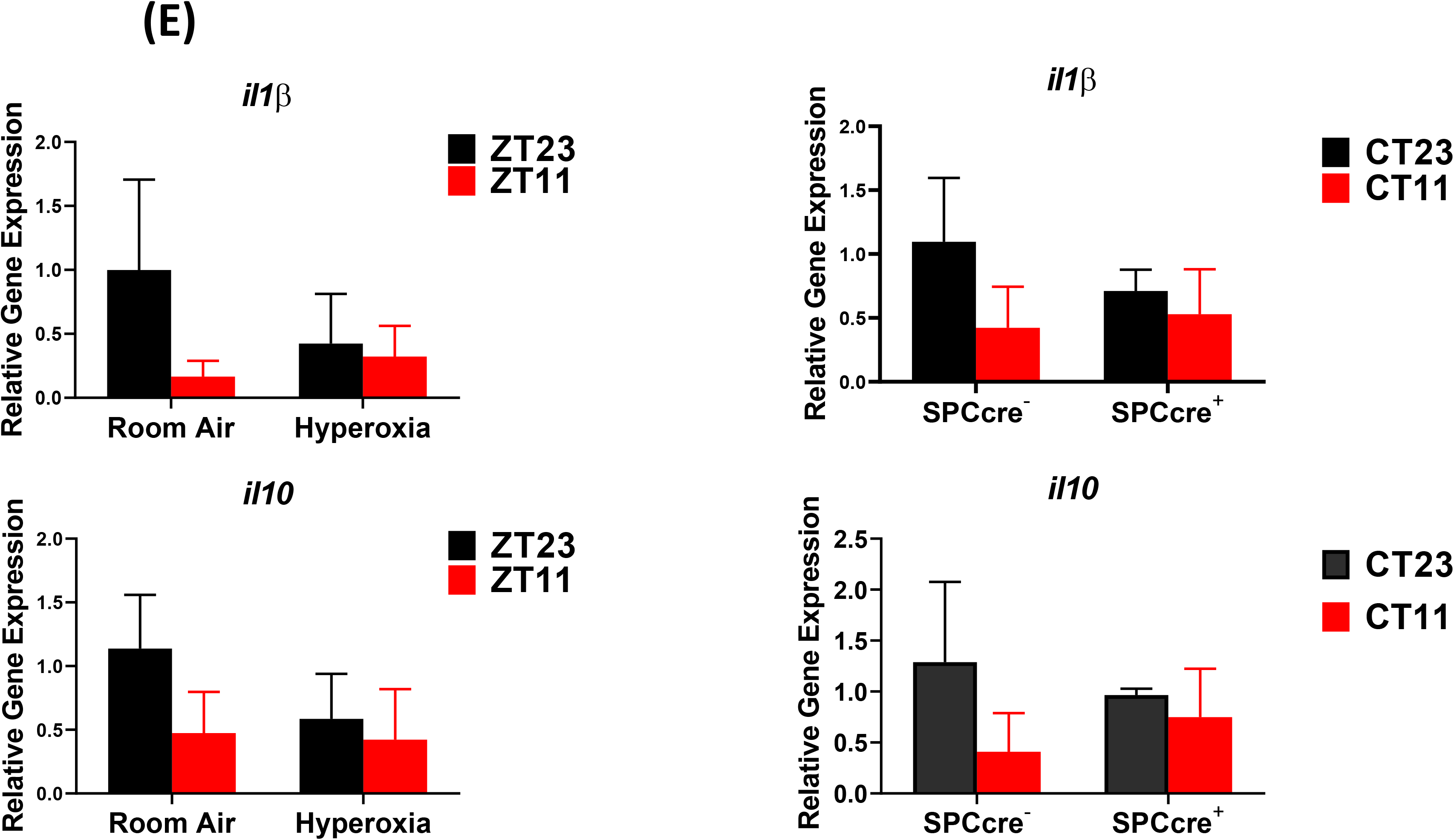
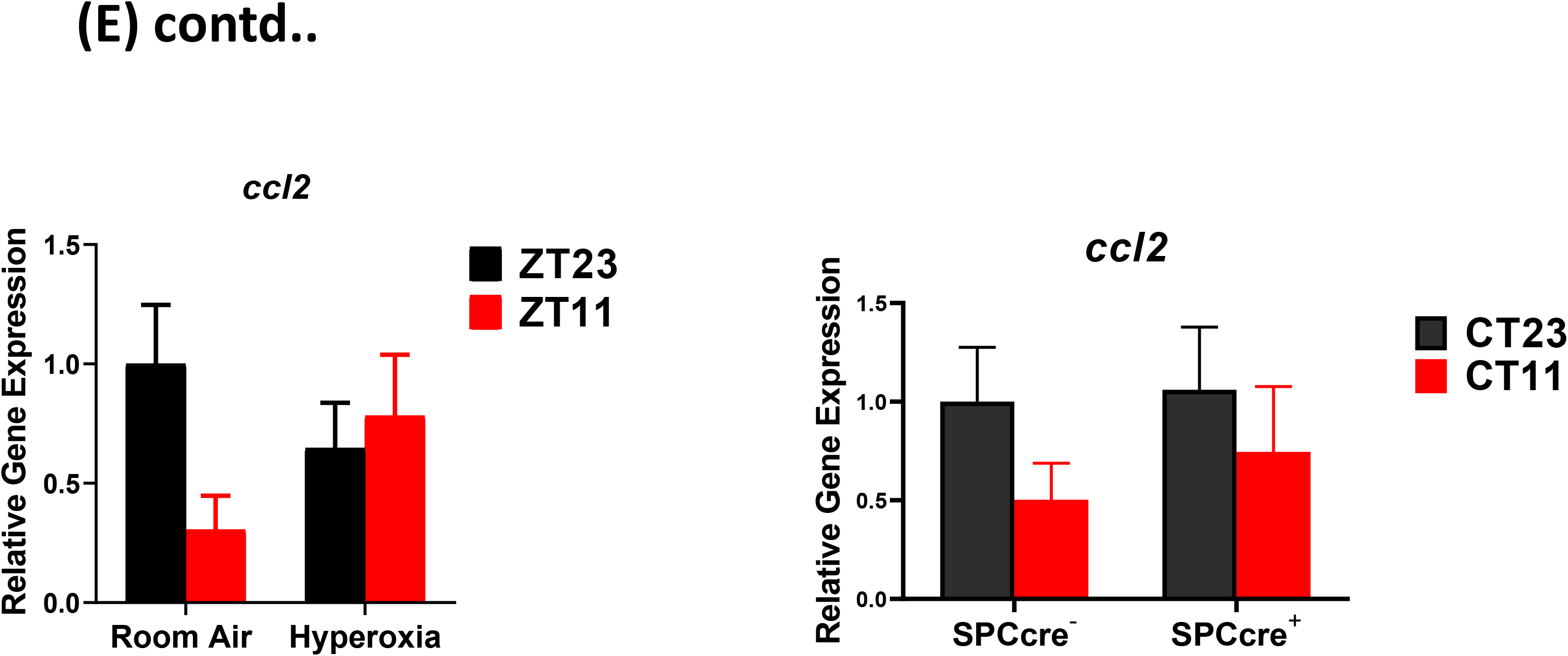

**Figure 6(F).**
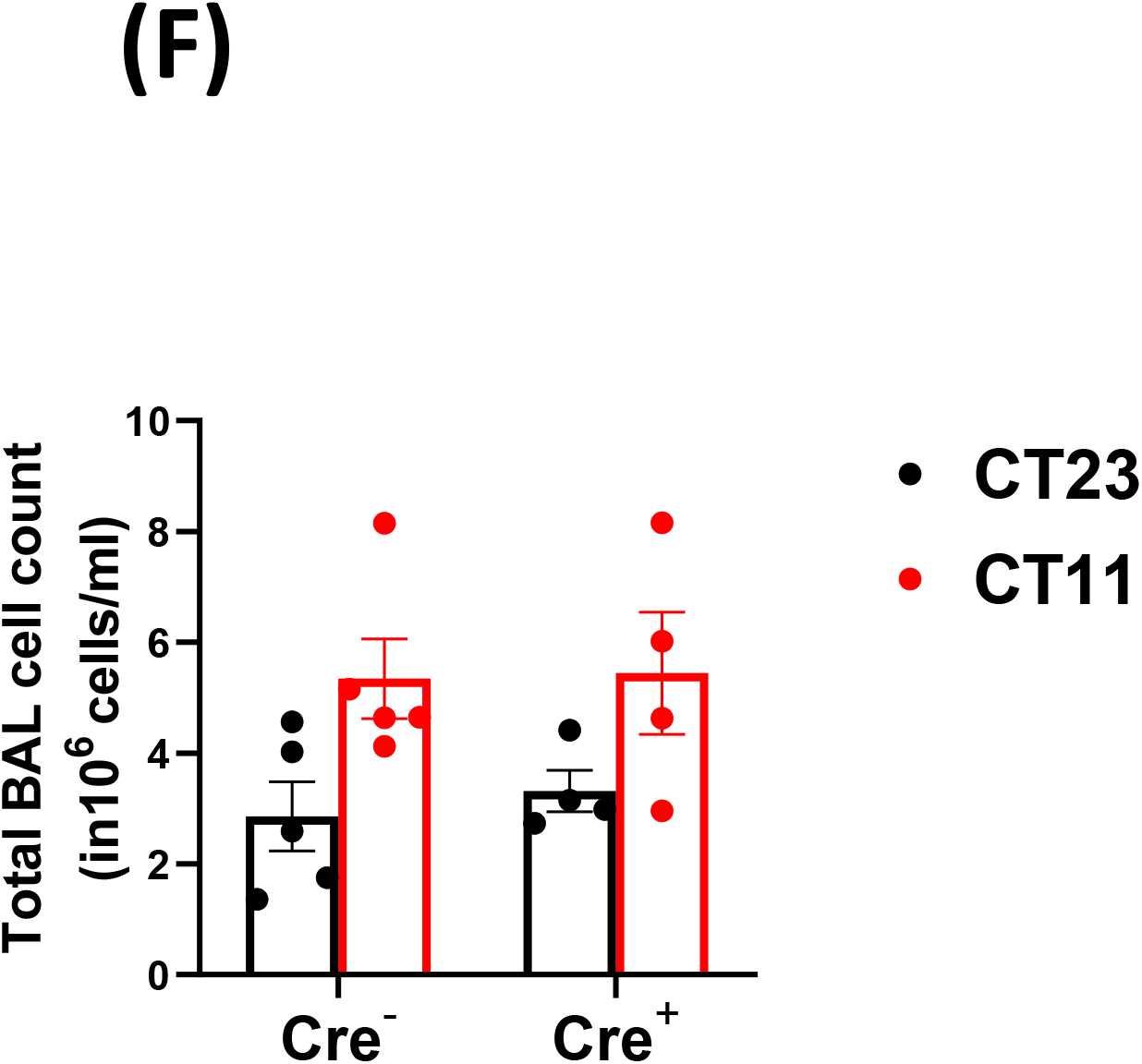

## Discussion

In the current study we systematically studied the effect of an early life hyperoxia exposure on the function of circadian rhythms in adulthood. A hallmark of circadian regulation is a difference in outcomes based on time of day at which the insult is sustained. We found that this time of day difference in outcomes from IAV is lost in adult animals exposed to hyperoxia as neonates, reminiscent of the effect of global genetic disruption of core clock gene Bmal1 from our previous work(13). Hyperoxia is known to cause significant negative effects in the lung and one speculation might be that the loss of the time of day difference in outcomes in reflective of a generalized lack of well-being rather than a loss of circadian control. However, our data clearly refutes that possibility, given that the hyperoxia exposed groups only ever have mortality comparable to the control group infected in the evening; never worse. We have also seen the loss of this circadian protection without worsening beyond the ZT11 group at lower doses of the virus (data not shown).

Consistent with our previous observations on clock disruption, the viral burden after challenge was similar across all the groups. Thus, hyperoxia does not affect viral replication: the loss of protection from IAV is not due to viral burden. Therefore, we addressed the possibility that this loss of central gating reflected dysregulation of central clock function (measured as locomotor activity rhythms), the immunological response to infection or peripheral clock function in pulmonary epithelial cells.

We found that mice exposure to neonatal hyperoxia did not have a significant effect on the central clock by adulthood. This is not surprising--while other adverse effects early in life are known to disrupt local or central circadian rhythmicity(21, 22), the SCN clock is well known to be resistant to non-photic stimuli(31, 32). At baseline, we also saw that neonatal hyperoxia would disrupt the immune clock, which would account for abrogated time of day protection from IAV. Since these cell populations were implicated in the exaggerated inflammation in the groups infected in the evening(13), we initially hypothesized that neonatal hyperoxia disrupts the immune clock which in turn abrogates the circadian protection from IAV. However, following IAV infection total BAL cell counts were higher in the group infected at ZT11 than in those infected at ZT23 in both the room air and neonatal hyperoxia exposed animals. This is in contrast with the work of O’Reilly et al who have shown that adults exposed to hyperoxia as neonates had a hyper-inflammatory response, with higher BAL counts in comparison to the room air controls. We speculate that this difference from the work by O’Reilly et al(10) results from our circadian experimental design. Further, we found that exposure to hyperoxia did not change the viral burden at either time points which is consistent with previous literature. Overall, given that even in our previous work, the myeloid clock did not completely control the phenotype seen in animals with a disrupted clock, we focused on the lung epithelium.

The lung consists of anatomically distinct regions (tracheas, bronchioles and alveoli) each populated by structurally and functionally distinct epithelial cell types. Tracheas and proximal airways are lined by multi-ciliated cells, secretory cells (Scgba1a1^+^), goblet cells and basal stem/progenitor (BSC) cells. The small airways terminate in the alveolar sacs which are mainly composed of alveolar type1 (AT1) or alveolar type 2 (AT2) cells(33). Although all epithelial cells are susceptible to IAV infection, AT2 cells are primarily affected by neonatal hyperoxia(27, 28). Even though there were only subtle differences in gene expression rhythms assayed in the entire lung, we could not exclude the possibility of changes in AT2 cells. We suspect that since AT2 cells contribute only a small portion of the total RNA extracted from the lung, the disruptive effect on the AT2 clock was underestimated. In fact, in our lung explant model while pulmonary rhythmicity was maintained, the amplitude was definitely blunted in the hyperoxia treated group. Thus, a role for the AT2 clock is supported by our *Bmal1^-/-^* data.

We and others(29, 34, 35) have demonstrated a role for the Scgb1a1 (or CCSP) clock in mediating lung injury, however, this is the first report of a cell intrinsic clock in AT2 cells. Although many clock genes participate in the generation and maintenance of circadian rhythms, *Bmal1* is the only circadian gene whose sole deletion is sufficient to cause arrhythmicity of locomotor activity—the hallmark of circadian disruption. Embryonic deletion of *Bmal1* results in a severe accelerated aging phenotype, in which the circadian phenotype is confounded by potentially off target developmental effects of Bmal1 deletion(30). We avoid any confounding by non-circadian effects from *Bmal1* by targeting gene deletion in adulthood. The time of day difference in IAV induced mortality in cre-littermates was abrogated amongst Sfptc-cre^+^ mice, treated with tamoxifen at 8 weeks of age. It is intriguing that the both hyperoxia and AT2 clock disruption not only abolish the circadian directed time of day protection from IAV, but also share many similarities in the mechanisms underlying this dysregulation. In both conditions, the loss of circadian regulation of the host response to IAV is not mediated through control of viral burden, but rather through worse injury. Overall, our data strongly support the possibility that hyperoxia disrupts the circadian regulation of lung injury in IAV through the AT2 clock. Since many of the Scgb1a1-cre models affect AT2 cells as well(36), it is possible that the Scgb1a1-cre described previously overestimates the effect of *bmal1* deletion. Thus, neonatal hyperoxia impairs the circadian regulation of host response to IAV in adult mice.

In conclusion, children born prematurely and suffering from even mild BPD have persistent adverse effects on their lung function into adulthood. This may result in part from long lasting damage to circadian mechanisms residing in AT2 cells that influence the response to injury after viral infection. The current study, provides a circadian paradigm for the long-term consequences of early life insults in this vulnerable population, paving the way for novel therapeutics and chronobiological strategies.

## Methods

### Animals, Hyperoxia exposure, Influenza infection

Newborn C57BL/6J mice (< 12hr old) were exposed to either 21% oxygen (room air) or ≥95% oxygen (O_2_) between postnatal days (PNDs) 0 and 5. Mice are born in the saccular phase of lung development and thus exposure to hyperoxia shortly after birth simulates the effects of hyperoxia on preterm lungs. This model of BPD is well established. Similar models have resulted in mild alveolar oversimplification and airway hyper-reactivity at 8 weeks of life(25). To minimize oxygen toxicity, nursing dams were switched out every 24 hours. Following exposure, the oxygen-exposed pups were recovered in room air with/alongside their littermates under 12hr LD conditions until 8-10 weeks of age.

For influenza infections, mice were acclimatized to reverse cycles in specially designed circadian chambers for 2 weeks. Thereafter they were infected at ZT23 or Z11 with 30 PFU of IAV (PR8) i.n. under light isoflurane anesthesia. This method has been used and validated in our previous work and is used by many circadian labs to remove confounding from infecting animals at different times of the conventional day(13).

#### Mouse Strains and Tamoxifen Administration

A mouse line with alveolar type II cell-specific knockout of *Bmal1* was generated by crossing *Sftpc-CreERT2*, a tamoxifen-inducible Cre, with *Bmal1^fl/fl^* mice. The *Sftpc-CreERT2* mice (on mixed Balbc/C567 background) were a kind gift from E.E. Morrissey (University of Pennsylvania; originally made in Dr. H Chapman’s group(37) and has been used widely in the field to target AT2 cells). The animals were back crossed for 4-5 generations onto a C57 background. Tamoxifen (Sigma-Aldrich) was dissolved/reconstituted in corn oil and ethanol to create a 100 mg/mL stock solution. To induce the cre driver, mice were given one dose of 5mg (in 50ul) of tamoxifen via oral gavage for 5 consecutive days around 8 weeks of life. Weights were monitored before, during and post-Tamoxifen administration and no significant weight loss was noted. They were acclimatized to reverse light-dark cycles for two weeks and exposed/placed in constant darkness 24-48 days before being infected with PR8 at (CT/ZT23 and 11).

### Flowcytometry

Lungs were harvested after perfusion with 10ml of PBS through the right ventricle. The lungs were digested using DNAse II (Roche) and Liberase (Roche) at 37°C for 30 mins. Dissociated lung tissue was passed through a 70 μm cell strainer, followed by centrifugation and RBC lysis. Cells were washed and re-suspended in PBS with 2% FBS. [(Details of the antibodies in table S1). 2-3 x 10^6^ cells were blocked with 1ug of anti-CD16/32 (Fc Block) antibody and were stained with indicated antibodies on ice for 20 minutes. No fixatives were used. Flowcytometric data was acquired using FACS Canto flow cytometer and analyzed using FlowJo software (Tree Star, Inc.). All cells were pre-gated on size as singlet live cells. All subsequent gating was on CD45+ in lung only. Neutrophils were identified as live, CD45^+^, Ly6G^+^ cells. Ly6C^hi^ monocytes were identified as live, CD45^+^Ly6G^−^Ly6C^hi^CD11b^+^ cells. NK cells were identified as CD45^+^Ly6G^−^ LysC^−^NK1.1^+^CD3^+^cells.

### Histology and Staining

Lungs from the mice were inflated and fixed with 10% neutral buffered Formalin (Sigma-Aldrich), embedded in paraffin, and sectioned. Lung sections were stained with hematoxylin-eosin. Stained slides were digitally scanned at 40x magnification using Aperio CS-O slide scanner (Leica Biosystems, Chicago IL). Representative images were taken from the scanned slides using Aperio ImageScope v 12.4. Soring was performed blindly using previously validated scoring method(13). Briefly the 8 lung fields selected at random 9at 20x magnification) were scored based on (1) Peri-Bronchial infiltrates (2) Per-vascular infiltrates (3) Alveolar exudates and (4) Epithelial damage of medium sized airways.

### Quantitative PCR

RNA was isolated/extracted from the inferior lobe of the mouse lung using TRIzol (Life Technologies). RNA was further purified using the RNeasy Mini Elute Clean Up Kit (Qiagen). The quantity and quality of RNA was assessed using the NanoDrop ND-1000 spectrophotometer (NanoDrop Technologies Inc). TaqMan and SYBR Green gene-expression assays were used to measure mRNA levels for genes of interest. Eukaryotic 18S rRNA (Life Technologies) and 28S (Sigma) were used as an internal control for TaqMan and SYBR Green assays, respectively. The samples were run on a LightCycler480 real-time PCR thermal cycler (Roche), and the relative ratio of the expression of each gene was calculated using the 2^ΔΔCt^ method. Gene expression values for each of *bmal1, cry1, per2, nr1d1* and *nr1d2* were fit by least squares to cosinor curves with a fixed period of 24 hours. An F-test comparing the full model allowing for differences in amplitude, phase and mesor between the hypoxia and room air conditions to the restricted model allowing only differences in mesor between conditions yielded no significant results at the p = 0.05 level.

### Running wheel activity

Mice were singly housed in circadian light control boxes (Actimetrics) to measure their locomotor activity, period, and amplitude using running wheels or Infrared Motion Sensors (Actimetrics). After the mice were acclimatized to circadian chambers, their activity was measured in 12-hour LD conditions for 2 weeks, constant dark conditions for 2-4 weeks, and again in 12-hour LD conditions. Data was analyzed using Clocklab software.

#### Ex Vivo bioluminescence recording from lung explants

lungs from Per2luc mice were harvested following perfusion to flush out blood cells. Thereafter, peripheral sections were obtained and explants were placed in media containing luciferin. Bioluminescence recording was initiated and data was recorded for 5-6 days. Data was analyzed using Clocklab software.

### Statistical analyses

Data was platted and analyzed using Graphpad Prism Software. Graphs represent data as means ± SE or median± IQR as applicable. Based on the distribution of data, either parametric (students t-test or ANOVA) or non-parametric (Mann-Whitney or Kruskal Wallace test) was used. Mortality data was analyzed using Mantel-Cox test. Bonferroni corrections was used for multiple comparisons.

#### Statement on rigor and reproducibility

All studies were performed using animals from either Jackson Labs and animals from in-house breeding. The background strain of each genetically modified animal has been specified and controls were cre-ve littermates on that same background. Reported findings are summarized results from 3-6 independent experiments.

### Approval

All animal studies were approved by the University of Pennsylvania animal care and use Committee and met the stipulations of the Guide for the care and Use of Laboratory animals.

## Acknowledgements

We are thankful to members of the FitzGerald lab-- Dr. S. Teegarden for help with animal breeding and general lab management; Dr. G. Grant for his comments and discussions at lab meetings. This work was supported by the NHLBI-K08HL132053 (SS), NICHD-K12HD043245 (SS), a Maturational Human Biology grant from the Institute of Translational Medicine and Therapeutics, University of Pennsylvania (SS), and NIH/NCRR RR023567 (GAF). Dr. FitzGerald is the McNeil Professor of Translational Medicine and Therapeutics and a senior advisor to Calico Laboratories.

## Author Contributions

SS and GAF conceived the project; YI, SS, AS and KNT designed experiments; YI, SS, SY, KNT and KF performed experiments and collected data. YI, AN, TB, NL and SS analyzed lung circadian gene expression data. YI wrote original draft with help from SS. GAF, AS, KNT and GSW helped with revisions of the draft. SS supervised all research activities.

