## Supplemental Figures for "Loss of Circadian Protection in Adults Exposed to Neonatal Hyperoxia"

#### Supplementary Figure 1

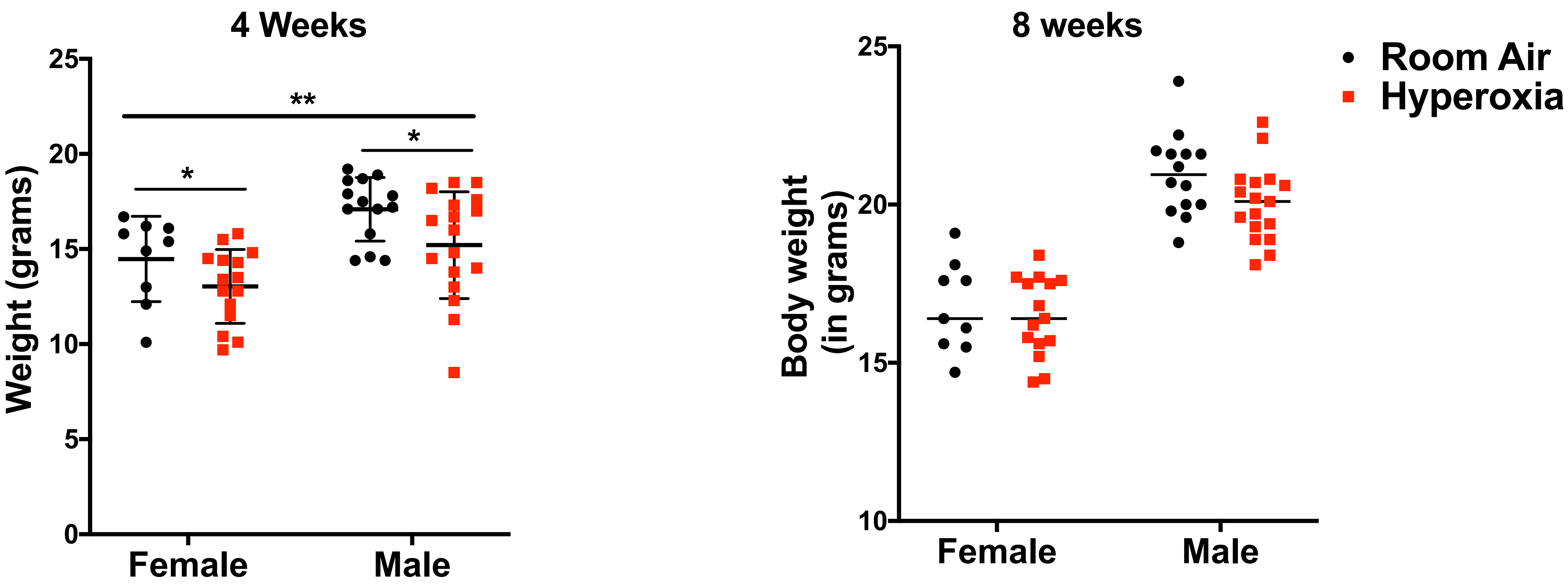

**Supplementary Figure 2**

Histology at 8 months of life

Hyperoxia/RA from $D_0$-$D_5$

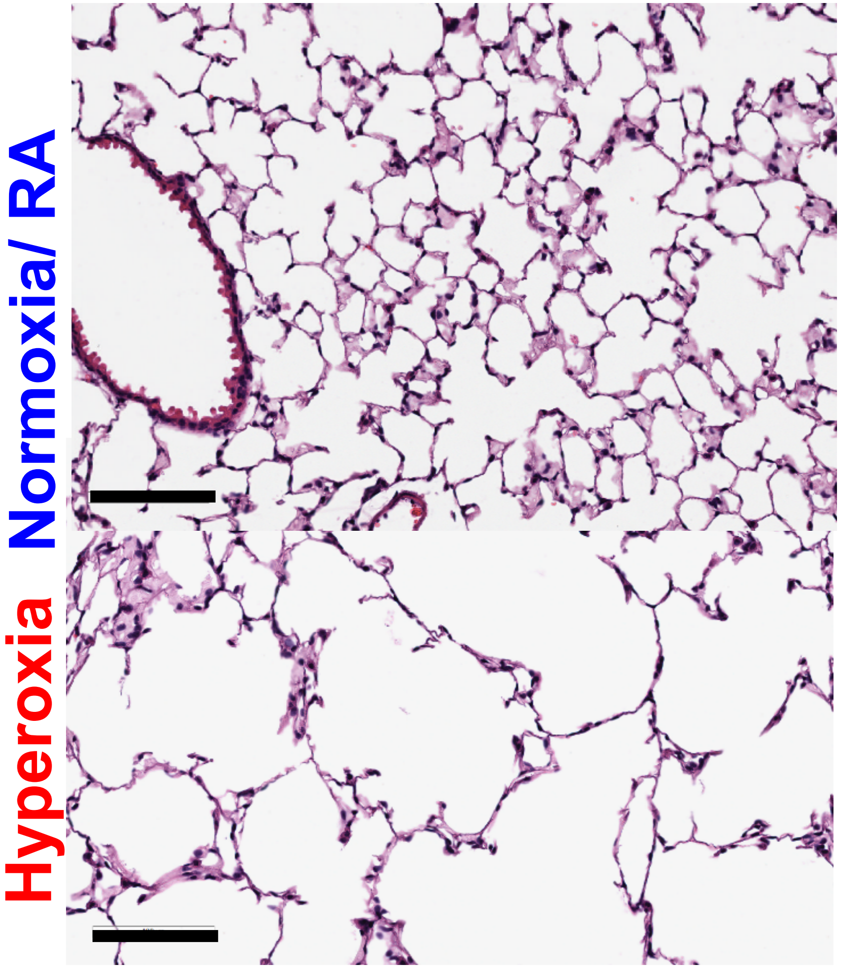

### Supplemental Figure 3

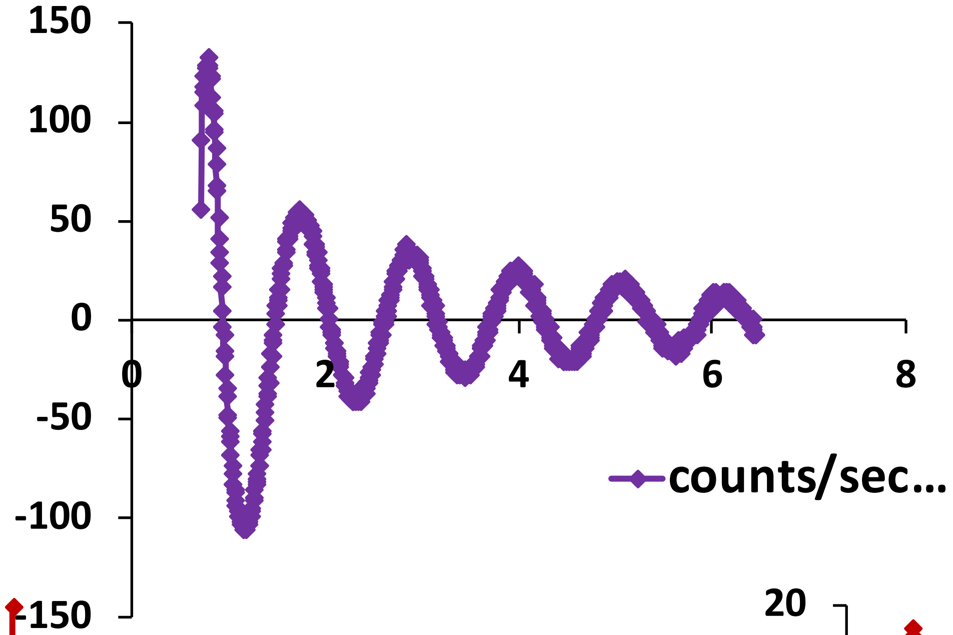


Exposed to ≥95% FiO2 for 5 days.
Rhythms assessed immediately
after exposure.

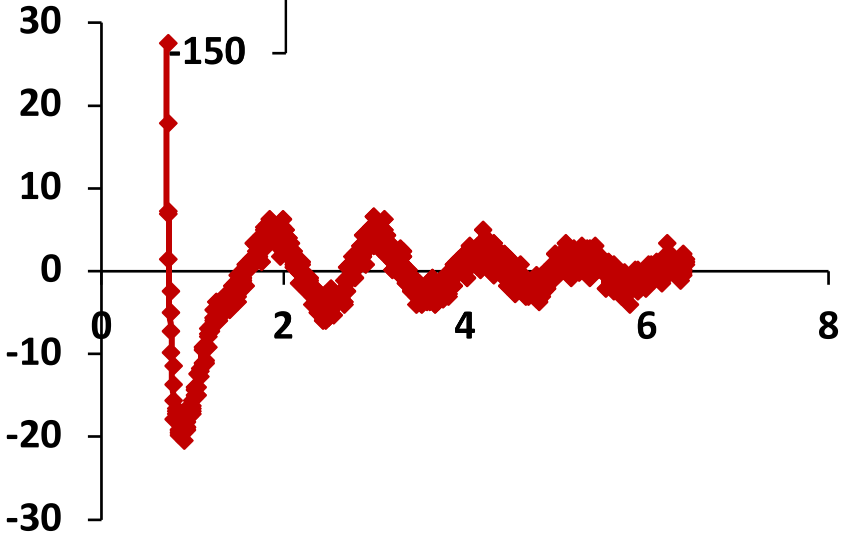


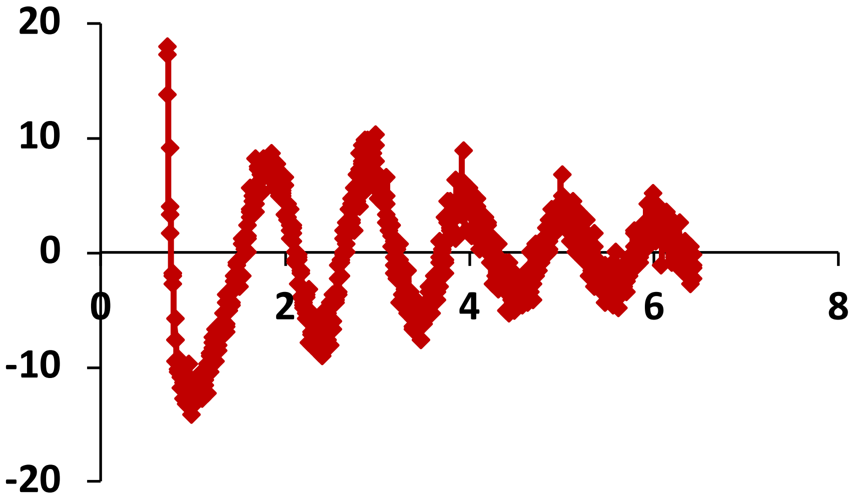

#### Legends: Supplemental Figures

**Figure S1. Body weights post hyperoxia exposure.**

After the 5-day neonatal hyperoxia exposure, body weight of both the hyperoxia exposed mice and their room air littermates were measured at 4 weeks and at 8 weeks as a measure of overall well-being (n= 9-15 per group for females and n= 16-17 per group for males).

**Figure S2.**

Representative micrographs of H&E stained lung sections taken from neonatal hyperoxia exposed mice and their room air littermates at 8 months of life. (photomicrograph bar=100μm).

**Figure S3.**

Representative bioluminescence tracing from the lungs of Per2luc mice immediately after exposure to 5 days of to ≥95% $FiO_2$. (n= 5 from 3 independent experiments)
